# Single Volume Image Generator and Deep Learning-based ASD Classification

**DOI:** 10.1101/763292

**Authors:** Md Rishad Ahmed, Yuan Zhang, Omer T. Inan, Hongen Liao

## Abstract

Autism spectrum disorder (ASD) is an intricate neuropsychiatric brain disorder characterized by social deficits and repetitive behaviors. Associated ASD biomarkers can be supportive of apprehending the underlying roots of the disease and lead the targeted diagnosis as well as treatment. Although deep learning approaches have been applied in functional magnetic resonance imaging (fMRI) based clinical or behavioral identification of ASD, most erstwhile models are inadequate in their capacity to exploit the data richness. Classification techniques often solely rely on region-based summary and/or functional connectivity analysis of one pipeline or unique site dataset. Besides these, biomedical data modeling to analyze big data related to ASD is still perplexing due to its complexity and heterogeneity. Single volume image consideration has not been previously investigated in classification purposes. By deeming these challenges, in this work, firstly, we design an image generator to generate single volume brain images from the whole-brain image of each subject. Secondly, the single volume images are analyzed by evaluating four deep learning approaches comprising one amended volume base Convolutional Neural Network framework to classify ASD and typical control participants. Thirdly, we propose a novel deep ensemble learning classifier using VGG16 as feature extractor to ensure further classification performance. Then, to evaluate the classifier performance across the inter sites, we apply the proposed method on each site individually and validate our findings by comparing literature reports. We showcase our approaches on large-scale multi-site brain imaging dataset (ABIDE) by considering four preprocessing pipelines, and the outcome demonstrates the state-of-the-art performance compared with the literature findings; hence, which are robust and consistent.

## I. INTRODUCTION

AUTISM Spectrum Disorder (ASD) is the disturbance of the structure and functioning of the brain that cause different abnormalities such as communication difficulties, social deficits, repetitive behaviors and cognitive delays, and nonsocial features such as restricted and stereotyped behaviors, all of which has a significant impact on adaptive functioning [1]–[3]. As investigated by the Centre for Disease, Control and Prevention in the United States, the estimated ASD prevalence of 1% or higher (1 subject in 59) and it is augmented dramatically in the last decades [4]. Therefore, finding precise biological markers to extrapolate the underlying roots of ASD pathologies and applying effective treatment for an individual are precarious.

One of the significant challenges in brain disorder research is to replicate the findings through larger datasets that can reflect the heterogeneity of clinical inhabitants. Functional magnetic resonance imaging (fMRI) has been extensively considered to perceive functional abnormalities of ASD patients which can characterize the neural pathways [5], [6]. Functional connectivity analysis has produced deep insights to see the brain abnormality connectomes between ASD and typical control (TC) either at individual or at group level characteristics. Recently, most of the machine learning techniques to study functional connectivity data rely on hand-engineered feature extraction, such as the correlation between region of interests (ROIs) and topological measurements of modularity, clustering based classification [7], segregation or integration [8]. On the other hand, brain ROIs which are the anatomical features or functional activations, and brain connectivity patterns, also a common approach to analysis ASD individuals based on the experts defined brain parcellation or data-driven strategy such as dictionary learning, clustering and ICA [9], [10]. Both the expert-defined and data-driven ROIs strategy have several complaints such as standardization, arbitrary decision, and selection of the regions that exhibit proficient information. A single volume image generator not only generates whole brain volume images but also ensure the coverage of the entire brain regions of each subject.

Machine learning (ML) such as SVM has been widely used to classify and exploit individual variation in functional connectivity of ASD [11]–[13]. Recently, deep learning models with neuroimaging modalities have been effective in identifying brain disorders such as ASD, Alzheimer Disease (AD) [14], [15]. With the rapid advancement of deep learning approaches for brain disorder diagnosis, convolutional neural networks (CNN) becomes the most popular method for ASD classification based on the functional connectivity or brain ROIs data [16], [17]. However, most deep learning approaches have been focused on functional connectivity or ROIs analysis, time-series data analysis, or temporal/spatial information of fMRI [18], [19] and also lack model transparency. The choice of the potential classification algorithm is another exception in the connectome-based analysis of ASD. Majority of the related works has been focused on simple linear predictive models using vectored connectivity correlation matrix. Additionally, ASD big data handling using deep learning techniques is still thought-provoking due to the lack of potential data mining and investigating methods from the heterogeneous, complex, and dynamic nature data to diagnose this brain disorder. Due to the heterogeneity, etiology, and severity of ASD, a more professed methodology is required to forecast and analyze the behavior and functionality of each subject. Hence motivated by these challenges discussed above, here we focus on designing a new image generator that can generate single volume images from whole-brain fMRI and propose two novel classification architectures for classifying ASD and typical controls. The single volume image portrays the brain regions in whole slices, conceding real-time subtraction of the acquiesced image, while ROIs only outline the interested brain regions depending on the slice assortment.

In this study, to the best of our knowledge for the first time, we design a single volume image generator that can produce 2D three-channel images from a functional magnetic resonance 4D NIFTI images. The main advantage of the single volume image generator is that, generated 2D images represent activated brain regions during the performed task by the patients promptly. Secondly, our key insight is to use these generated images as the input of four deep learning approaches, including an improved convolutional neural network, to classify ASD and typical controls. Thirdly, different from the traditional loss function, we derive a combined loss function based on the weighted class wise loss and regularization for reliable feature extraction and classification. Fourthly, we propose a novel deep ensemble learning framework based on the VGG16 as a feature extractor from the generated images to ensure further classification performance. Finally, to evaluate the classifier performance and check the data variability across the sites, we apply the proposed method on each site individually and validate our findings by comparing literature reports. The proposed approaches with a combined loss function and generated single volume images, establishes a novel benchmark model for ASD detection on ABIDE database.

The leftover of the paper is organized as follows: Section 2 discusses the related works; Section 3 covers broad methodological explanation including single volume image generator and propose deep learning models. In section 4, the Experiment and Discussion of this method including dataset are presented and finally Conclusion is drawn in Section 5.

## II. RELATED WORKS

The amalgamation of machine learning methods and brain imaging data permit the classification of ASD, which can assuage the significant suffering and provide safety for this neuropsychiatric brain disorder. Studies on ASD classification using different imaging modalities and their analysis approaches, specifically deep learning techniques, are discussed in this section.

Study on the functional connectivity of brain networks is a sturdy utensil to understand the neurological bases of a diversity of brain disorders such as autism [20], [21]. In the work [22], Abraham et al. used resting-state functional MRI to extract functionally-defined brain areas and support vector classifier (SVC) to compare connectivity between ASD and typical control. They used 871 subjects from ABIDE dataset for connectome-based prediction and got 67% accuracy. Based on the patterns of functional brain connectivity, Heinsfeld et al. used denoising auto-encoder to classify and unveil the neural patterns of ASD [23]. They achieved an accuracy of 70% on CPAC pipeline, and the pattern that emerged from the classification presented an anticorrelation of brain function between anterior and posterior areas of the brain. Deep learning-based feature extraction method has been used by Choi to identify multivariate and nonlinear functional connectivity patterns of ASD using DPARSF pipeline data in [24]. The predefined 90 brain regions were used to generate connectivity matrix, and variational auto-encoder (VAE) generator was used to summarize the connectivity matrix. Guo et al. were used multiple stacked auto-encoder (SAE) as a feature selection method from whole-brain FCP obtained by Pearson correlation of ROIs [25]. Using only one data site (UM), they got a classification accuracy of 86.36%. In [26], using only CCS pipeline without global signal regression data and LSTMs method for classification of individuals with ASD, Dvornek et al. achieved 68.5% accuracy.

On the other hand, time-series for several sets of regions-of-interests (ROIs) also have the potentiality to classify and see brain network connections of ASD. ROIs are usually computed using a predefined atlas or a parcellation scheme on anatomical features, functional activations, and connectivity patterns of brain [10], [27]. Dvornek et al. [28], has been incorporated phenotypic data with rsfMR into a single LSTM based model for classifying ASD and achieved an accuracy of 70.1%. They used CCS pipeline data without global signal regression and cross-validation framework. With the development of deep learning model specifically, Convolutional Neural Networks (CNNs) have found abundant applications on 2D and 3D images which can exploit image intensities and pixel grid to decipher image segmentation and classification problems [29]. Khosla et al. used stochastic parcellation and seven atlases to classify ASD based on the 3D CNN approach [30]. The authors also performed an ensemble learning strategy corresponding to different brain parcellation for combining predictions. Ktena et al. highlighted the potential of convolutional neural network models for connectome-based classification. Graph convolutional neural networks (G-CNN) [31], is another way to identify brain patterns that can act as a neuropathological biomarker. Using CPAC preprocessing pipeline and atlases (Harvard-Oxford) in [32], Anirudh et al., achieved 70.86% classification accuracy based on the G-CNN and ensemble learning.

From the analysis of the recent research work on ASD, we notice that most of the researcher used functional connectivity (FC) or ROIs data for classification purposes. Besides, the majority work belongs to either one pipeline or one site and one atlas images for classification, which is not plentiful to answer some research and clinical questions. Biomedical big data is generally preferable to improve the dependability and core contributions of research for the treatment of brain disorder like ASD. By considering these challenges, we design an image generator to generate the single volume brain images from the whole brain image using four preprocessing pipelines mentioned by ABIDE. Furthermore, we observe that extracting features from the images using deep learning models also enhance classification performance. We also propose a novel deep ensemble learning model which can be treated as a new benchmark approach for ASD classification compared with the recent literature reviews.

## III. METHODOLOGY

### A. Data Preprocessing Pipelines

The ABIDE data instigated by the various preprocessing pipelines are analogous, and there is no consensus on the superlative methods [33]. The foremost difference between preprocessing pipelines are the precise algorithms and parameters used for each of the steps and software implementations. Rather than being doctrinaire and preferring a single strategy, we analyzed four different preprocessing approaches. The principal advantages of four strategies that, it will overcome the controversies surrounding by bandpass filtering and global regression. Selected strategies are Connectome Computation System (CCS), Configurable Pipeline for the Analysis of Connectomes (CPAC), Data Processing Assistant for Resting-State fMRI (DPARSF) and Neuroimaging Analysis Kit (NIAK). Table I arranges for a summary of the distinct preprocessing steps and how they vary across pipelines. As mentioned by ABIDE, functional processing was performed using only four strategies which we covered in this study; however other strategies were used for structural preprocessing and calculation of cortical measures.

**Table I:** Parameters and steps overview of four different functional preprocessing stratagems.

|  | Steps | CCS | CPAC | DPARSF | NIAK |
| --- | --- | --- | --- | --- | --- |
| Basic Processing | Slice timing Correction | Yes | Yes | Yes | No |
|  | Motion Re-alignment | Yes | Yes | Yes | Yes |
|  | Intensity Normalization | Yes | Yes | No | Non-uniformity |
| Nuisance Signal | Motion | 24-param | 24-param | 24-param | scrubbing and 1st principal component |
| Removal (Regressor) | Tissue signals | mean WM and CSF signals | CompCor (5 PCs) | mean WM and CSF signals | mean WM and CSF signals |
|  | Motion re-alignment | Yes | Yes | Yes | Yes |
|  | Low-frequency drifts | linear and quadratic trends | linear and quadratic trends | linear and quadratic trends | discrete cosine basis with a 0.01 Hz high-pass cut-off |

In this work, data are selected from the filt_global preprocessing stratagem, which is band-pass filtered (0.01-0.1 Hz) and spatially registered using a nonlinear method to MNI152 template space for each of four pipelines.

### B. Single Volume Image Generator

Functional MRI data is a set of volume images that are assimilated over a repeated period. So there needs a tool to generate the images between task and rest periods for scrutinizing the activated brain regions. Rather than traditional analysis of functional connectivity or brain ROIs, in this work, we design an image generator to produce single volume brain images from the preprocessed whole-brain functional image. Single volume image depicts the 2D visualization of the brain activity by considering each time interval of each voxel. Fig. 1 represents the flow diagram to generate the single volume images by predefined displaying mode. The working principle of the single volume image generator has several folds: firstly, the generator checks out the shape of each input Nifti files, the input shape might be the 4D fMRI images. The generator main body, which is the conditions to draw output depending on the enumeration and iteration counter inside the voxel image. Then set the corresponding parameters to demonstrate the real-time brain activations. Finally, plot and save the brain images by counting the whole brain volumes into two types of volumetric images for each ASD and TC individuals. The detailed description of the plotted two types of volumetric images is explained below.

**Figure 1:**
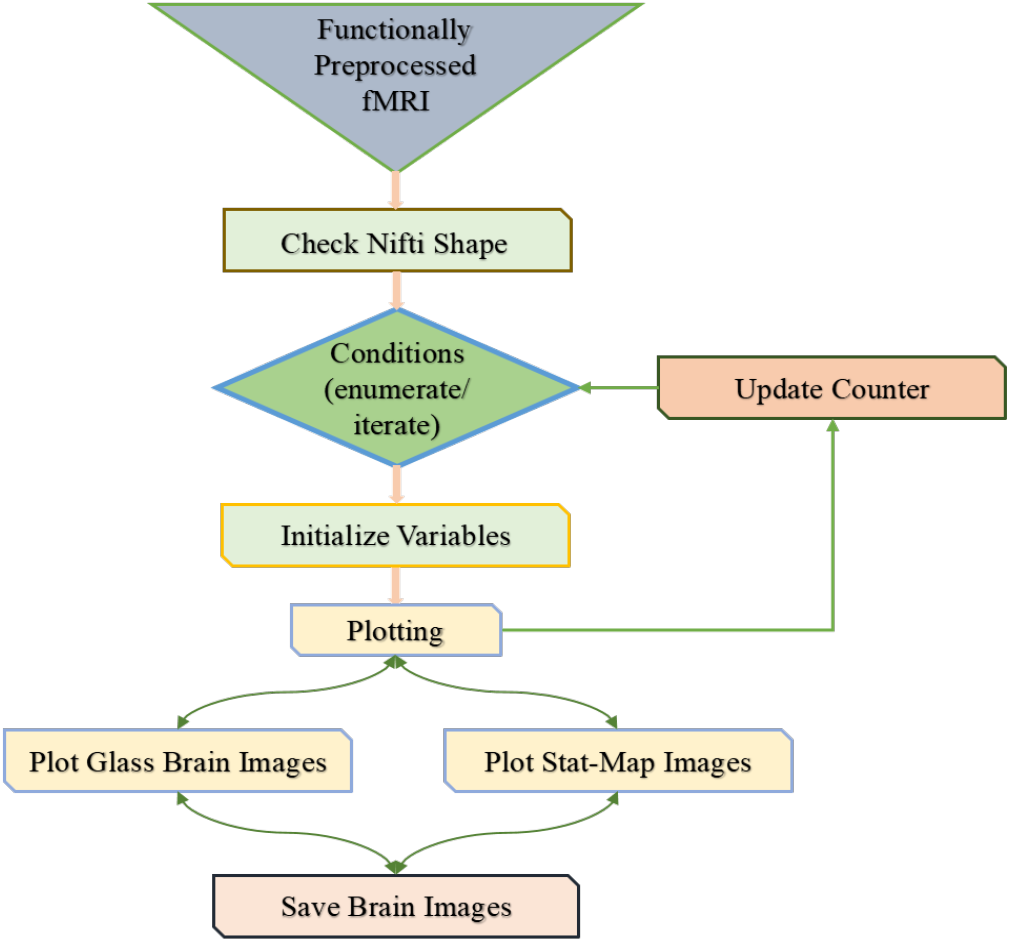
The flow diagram to generate single volume glass brain images.

#### Glass Brain and Stat_Map Images

The glass brain is a 3D brain visualization that displays real-time source activity and connectivity between brain areas [34]. On the other hand, Stat_Map is the full name of the statistical images which plot cuts of an ROI/mask image. We prefer glass brain and stat_map displaying mode for a single volume image because of its power of projecting high-resolution 3D model of an individual’s brain, skull, and scalp tissue. The corresponding parameters to display and save the single volume images from 4D multiple brain volumes image by using the proposed single volume image generator are shown in Table II. The plotted images were in MNI space for all the considered pipelines to work image function accurately.

**Table II:**
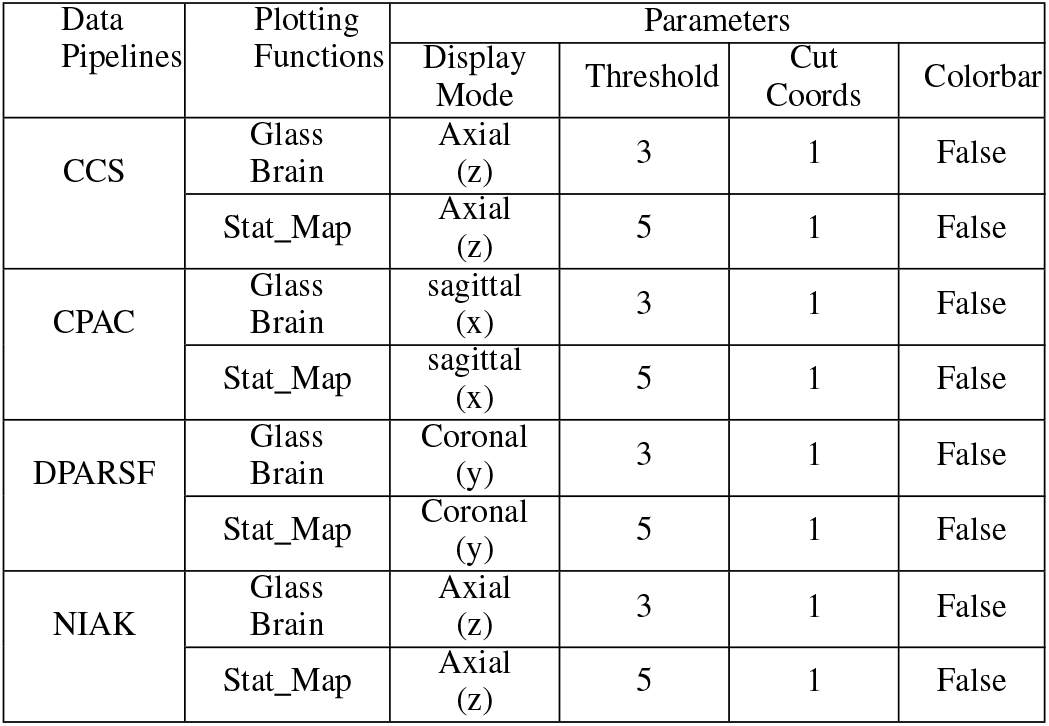
The corresponding parameters to display and save images by using the proposed single volume image generator.

| Data Pipelines | Plotting Functions | Parameters |  |  |  |
| --- | --- | --- | --- | --- | --- |
|  |  | Display Mode | Threshold | Cut Coords | Colorbar |
| CCS | Glass Brain | Axial (z) | 3 | 1 | False |
|  | Stat_Map | Axial (z) | 5 | 1 | False |
| CPAC | Glass Brain | sagittal (x) | 3 | 1 | False |
|  | Stat_Map | sagittal (x) | 5 | 1 | False |
| DPARSF | Glass Brain | Coronal (y) | 3 | 1 | False |
|  | Stat_Map | Coronal (y) | 5 | 1 | False |
| NIAK | Glass Brain | Axial (z) | 3 | 1 | False |
|  | Stat_Map | Axial (z) | 5 | 1 | False |

### C. Proposed Deep Learning Architecture

Fig 2 depicts the overall deep learning architecture specifically based on CNN for classification of ASD. Convolutional neural network, as a portion of the neural network is widespread for task interrelated to image classification and segmentation. CNN debilitates the limitations of traditional neural networks through a local connection, sharing weights and sampling. There exist two rudimentary operations in CNN: convolution through a kernel (weights and bias to convolve input feature map) and subsequent sampling of the convolved feature map.

**Figure 2:**
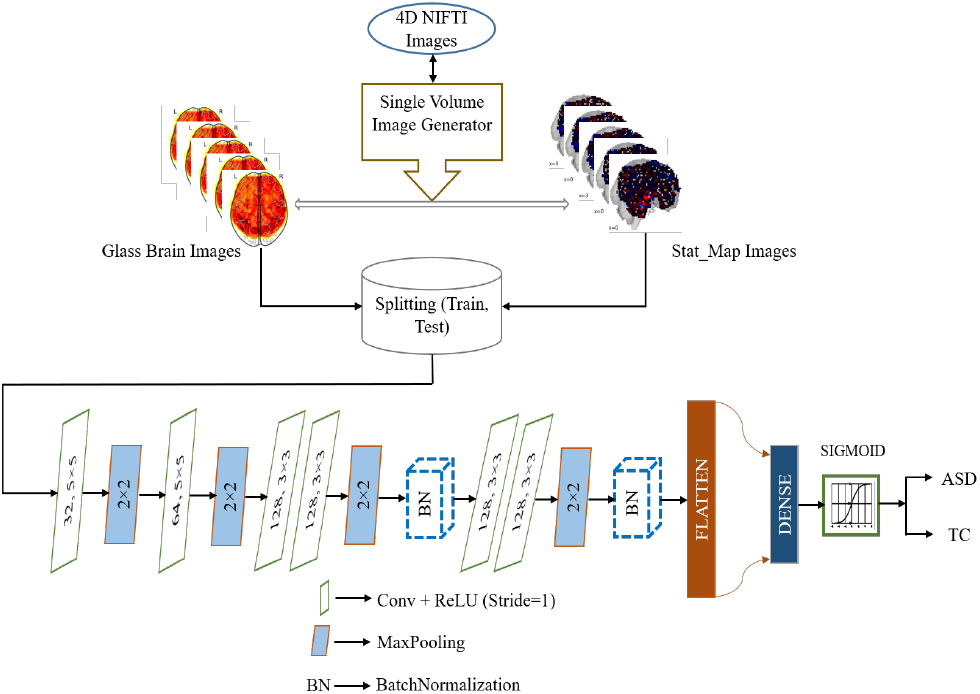
Proposed CNN architecture for classifying ASD vs. TC

From Fig. 2, the enhanced CNN architecture has six convolutional layers, four max-pooling, two batch normalization layer, one flatten and dropout layer with ratio 0.25, and two densely connected layers followed by a sigmoid activation function. The number of kernels and layer types are signified in each box. To conciliate the model slow training due to the layer parameter changes, we introduce the batch normalization [35] layer, which provides much higher learning rates and flout initialization.

#### Loss Function

We contemplate the class-wise crossentropy loss as given below:

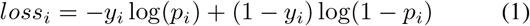

After that, we measure the higher weight (*w*_*i*_) to give it in losses incurred on the training data samples.

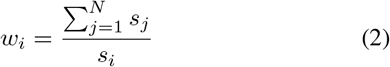

where *s_i_* is the number of training samples per classes. Hence the weighted binary cross entropy loss can be calculated from *equation* (1) a*nd* (2) as,

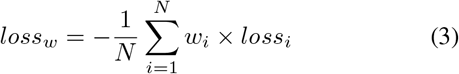

To penalize the sum of the absolute value of weights and solve the overfitting problem, we added *L*_1_ regularization, as defined below.

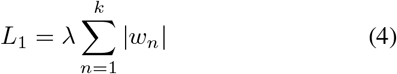

We define a new loss function by combining weighted loss with regularization. The combined loss function is given by using *equation* (3) a*nd* (4) as,

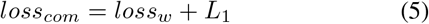

#### Benchmark CNN Approaches

To analyze our generated single volume images, we executed Keras based pretrained benchmark CNN approaches as follows: DenseNet [36], ResNet [37], Xception [38], and Inception V3 [39]. For classification of ASD, the output node-set to a sigmoid function, and binary cross-entropy loss was used. The models are trained with a mini-batch size of 64, ADAM was used as the optimizer with learning rate 0.0004, *beta*_1 = 0.9, and *beta*_2 = 0.999.

### D. Proposed Deep Ensemble Classifier

Ensemble learning framework sometime overcomes the limitations of traditional deep learning models which often rely on ROIs based summary statistics and linear models for ASD classification [30]. In our experiments, we reconnoitered four ensemble learning strategies by combining benchmark approaches with improved CNN model. Each ensemble model was trained and tested using two types of generated images from each pipeline. Fig 3 shows the proposed deep ensemble learning architectural overview for ASD classification. Here, we used VGG16 [40] model to extract features from both generated images. Each ensemble classifier averages the predictions of the models for a specific pipeline using each one of the two types of images. The final evaluation metrics are calculated as the two-stage arithmetic mean of the specific binary class predictions for classification. The mathematics behind this is given below:

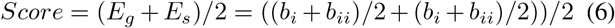

where *E_g_* and *E_s_* denotes the glass brain and stat_map image ensemble classifier respectively and, *b_i_* and *b_ii_* are the base classifier.

**Figure 3:**
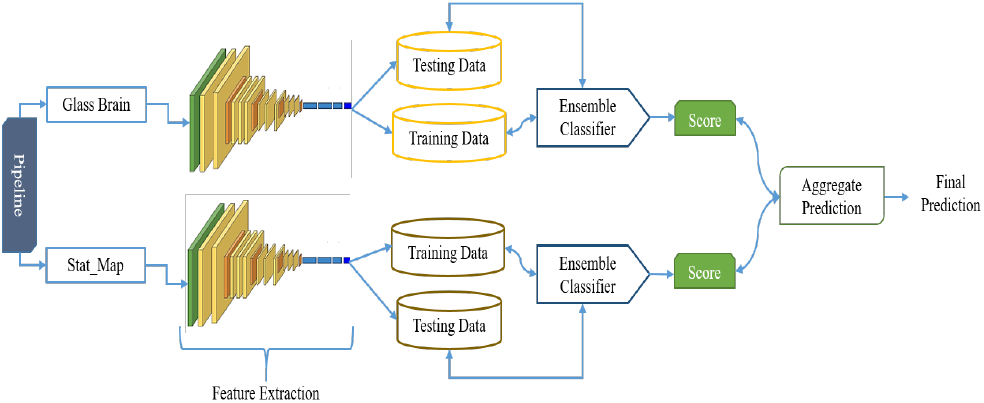
Architectural overview of deep ensemble learning for ASD classification.

The activation of the last layer was set to sigmoid function for the classification task. Each ensemble model was trained for 1000 iterations using ADAM optimizer with learning rate 0.0004, *beta*_1 = 0.9, and *beta*_2 = 0.999.

## IV. EXPERIMENTS AND DISCUSSION

### A. Participants

Resting-state fMRI allows for the investigation of the disturbance of brain networks without the added complexity of variation associated with task-related brain activation. The present study was carried out using fMRI data from the ABIDE1, which is a particularly complex dataset due to its heterogeneity, vast range subjects comprised and different imaging protocols. It’s also a connotation that provides previously collected multi-site ASD and matched typical control data in the scientific research community [33]. One of the fundamental benefits of ABIDE is that they have available preprocessed data using different preprocessing pipelines. The fMRI preprocessed data was downloaded through four preprocessing pipelines from the Preprocessed Connectomes Projects^1^. We incorporated data from 529 ASD individuals and 573 typical controls (TC) for each pipeline, which is comprised of 17 different imaging sites. Table III contains the key phenotypical information, including distribution of ASD and TC by sex and age, and the ADOS score, where Ψ means that the site did not have this information.

**Table III:** Phenotypic summary of ABIDE-I functional Dataset.

| Site | Count |  | Sub. Total | Age Avg.(SD) | ADOS- ASD(SD) |
| --- | --- | --- | --- | --- | --- |
|  | ASD | TC |  |  |  |
| Caltech | 19 | 19 | 38 | 28.16 (10.6) | 13.1 (4.5) |
| CMU | 14 | 13 | 27 | 26.6 (5.7) | 13.07 (3.12) |
| KKI | 22 | 33 | 55 | 10.09 (1.33) | 12.55 (3.53) |
| Leuven-1 | 14 | 15 | 29 | 22.59 (3.55) | $\Psi$ |
| Leuven-2 | 15 | 20 | 35 | 14.16 (1.42) | $\Psi$ |
| MaxMun | 24 | 33 | 57 | 26.16 (12.08) | 9.5 (3.56) |
| NYU | 78 | 106 | 184 | 15.25 (6.58) | 11.30 (4.08) |
| OHSU | 13 | 15 | 28 | 10.81 (1.87) | 9.69 (3.54) |
| Olin | 20 | 16 | 36 | 16.81 (3.49) | 14.1 (4.04) |
| Pitt | 30 | 27 | 57 | 18.90 (6.88) | 12.38 (3.22) |
| SBL | 15 | 15 | 30 | 34.37 (8.60) | 9.25 (1.67) |
| SDSU | 14 | 22 | 36 | 14.41 (1.84) | 11.15 (4.26) |
| Stanford | 20 | 20 | 40 | 9.96 (1.58) | 11.74 (3.23) |
| Trinity | 24 | 25 | 49 | 17.18 (3.64) | 10.54 (2.87) |
| UCLA_1 | 41 | 32 | 73 | 13.16 (2.40) | 10.24 (3.56) |
| UCLA_2 | 12 | 14 | 26 | 12.49 (1.53) | 13.25 (2.67) |
| UM_1 | 55 | 55 | 110 | 13.40 (2.89) | $\Psi$ |
| UM_2 | 13 | 22 | 35 | 15.96 (3.32) | $\Psi$ |
| USM | 58 | 43 | 101 | 22.10 (7.68) | 9.03 (6.45) |
| Yale | 28 | 28 | 56 | 12.71 (2.88) | 11 ( $\Psi$ ) |

### B. Visualization of the Generated Single Volume Images

The envisioned image generator as discussed previously, firstly ensures the shape of the functionally preprocessed images. If the checked image shape is 4D NIFTI (.nii.gz), then display the brain images according to the volume of the fMRI image with the set parameters. For instance, if an image has volume 176, the generator, therefore generates 176 brain images. Finally, the generator automatically saves the single volume images from the exposed whole volume brain images in PNG format according to the defined label. The produced glass brain and stat_map images for the first ten subjects of CCS, CPAC, DPASRF, and NIAK pipelines from both ASD and TC are shown in Fig 4, Fig 5, Fig 6 and Fig 7, respectively. In Fig 4–7, a and b represents the corresponding glass brain and stat_map images of ASD and TC individuals, respectively.

**Figure 4:**
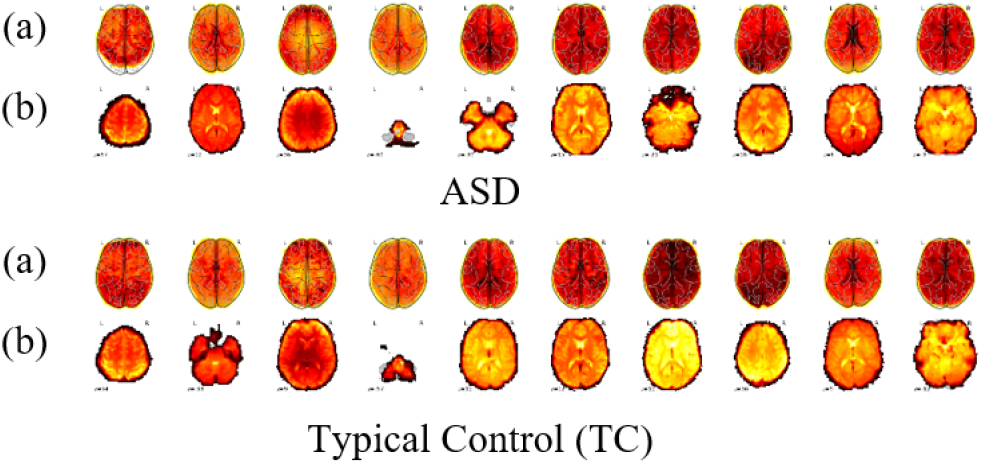
Axial view (z) of the CCS pipeline images. .

**Figure 5:**
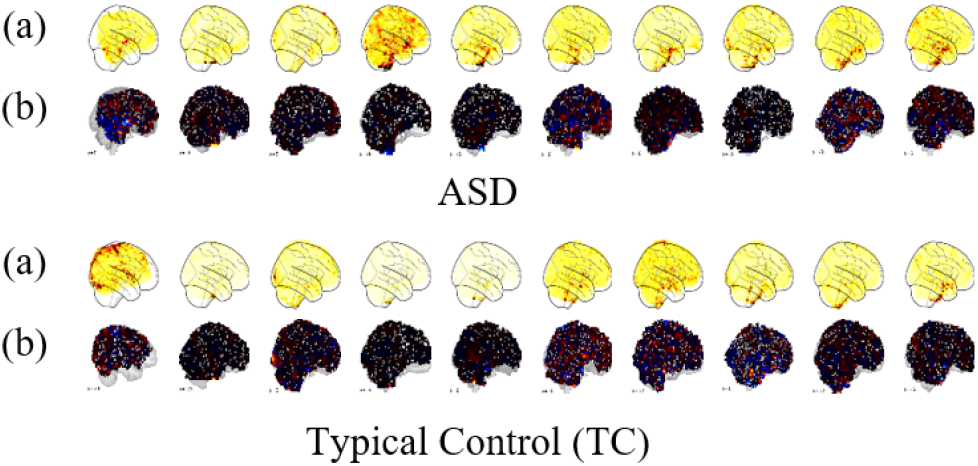
sagittal view (x) of the CPAC pipeline images.

**Figure 6:**
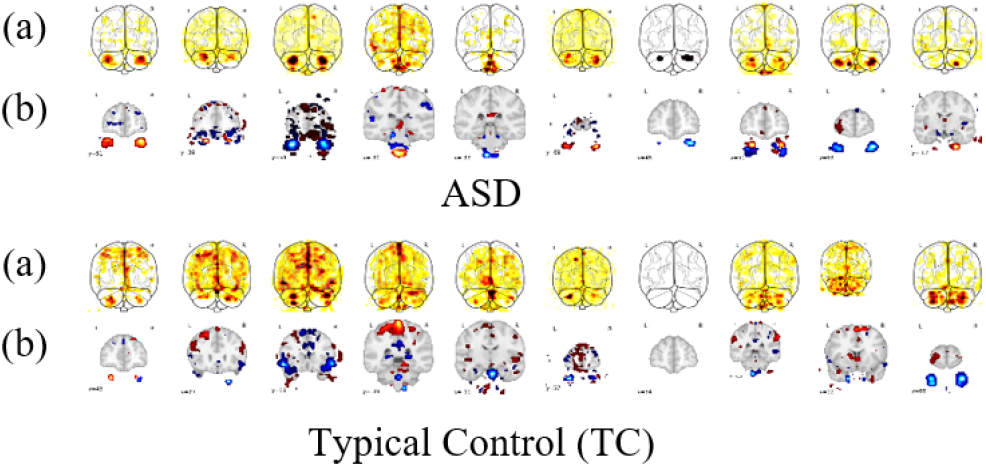
Coronal view (y) of the DPASRF pipeline images.

**Figure 7:**
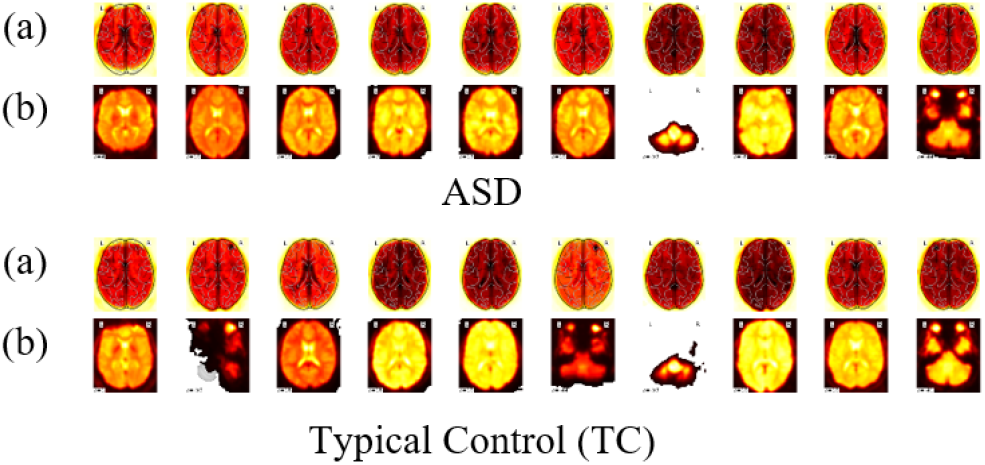
Axial view (z) of the NIAK pipeline images.

From the analysis of the visualized images, Fig 4 and Fig 7 shows the horizontal slice that divides the brain into upper (superior) and lower (inferior) segments and is an x-z plane. On the other hand, Fig 5 illustrates the longitudinal slice that separates the brain into left and right halves and is also an x-z plane. It is called the sagittal view because it transpires through the sagittal stitch. From Fig 6, it depicts the frontal spheres that divide the brain into the front (posterior) and back (anterior) parts and is an x-y plane.

Finally, we processed the whole dataset according to the model requirements because of the shape of the generated images varies with different volume for different imaging sites as well as parameter selection during image acquisition.

### C. Model Training and Testing

In our experiment, to test the robustness of the generated images and improved model, we first divided the whole dataset into two parts: training (85%), and testing (15%). From the generated training images, approximately 85% of the whole images were considered as the training set, and 15% as the validation set. We reckon every image according to their subject’s image as it is either ASD or TC for binary classification. During training the first model (Fig 2), we considered Keras based ImageDataGenerator to process the generated images according to model input requirements. By setting up the necessary parameters such as batch size (64), rescale (1. /255), class mode (binary), we reshaped the images in target size (160,160) of RGB images. On the other hand, in ensemble learning classifier (Fig 3), we incorporated VGG16 model for feature extraction and then split the data into two parts training and testing to feed into the classifier models.

### D. Classification Performances Evaluation on ABIDE Dataset

Deep learning approach and ABIDE data have previously been studied to identify and analyze of ASD showing different measurement metrics. In this work, we evaluate four performance measurement metrics such as precision (P), recall (R), F1-score (F) and accuracy (A) to validate the algorithm performance. Precision is outlined as the ratio of the correctly ASD positive labeled to all ASD positive labeled, and recall is the ratio with the whole ASD positive in reality. On the other hand, F1-score considers both precision and recall measurement. Accuracy is delineated as the ratio of correctly labeled individuals to the whole number of subjects. All of the four metrics are used to validate the classification ability of our model for ASD and TC classification. Table IV shows the comparison between different performance measurements for benchmark and proposed method using two categorised images of four pipelines.

**Table IV:** Corresponding precision (P), recall (R), F1-score (F) and accuracy (A) comparison of ASD classification.

| Pipelines | Generated Image | DenseNet (%) |  |  |  | ResNet (%) |  |  |  | Xception (%) |  |  |  | Inception V3 (%) |  |  |  | Proposed Method (%) |  |  |  |
| --- | --- | --- | --- | --- | --- | --- | --- | --- | --- | --- | --- | --- | --- | --- | --- | --- | --- | --- | --- | --- | --- |
|  |  | P | R | F | A | P | R | F | A | P | R | F | A | P | R | F | A | P | R | F | A |
| CCS | Glass Brain | 79.1 | 78.2 | 78.6 | 80.1 | 78.5 | 79.2 | 78.8 | 79.8 | 80.4 | 77.8 | 79.1 | 82.4 | 80.1 | 78.9 | 79.5 | 80.1 | 77 | 79.5 | 78.3 | 80.2 |
|  | Stat_Map | 78.2 | 77.8 | 78 | 79.8 | 78.3 | 80.3 | 79.3 | 81.9 | 77.6 | 79.6 | 78.6 | 80.4 | 78.1 | 79.2 | 78.6 | 79.3 | **80.5** | 79.3 | 79.9 | 79.3 |
| CPAC | Glass Brain | 78.9 | 76.9 | 77.9 | 79 | 79.4 | 78.4 | 78.9 | 79.7 | 78.6 | 76.9 | 77.7 | 82.6 | 78.8 | 77.2 | 78 | 80.8 | 78.2 | 79.5 | 78.9 | **83** |
|  | Stat_Map | 77.9 | 79 | 78.5 | 80 | 76.9 | 78.8 | 77.8 | 80 | 79.3 | 78.9 | 79.1 | 79.5 | 78.8 | 77.9 | 78.3 | 79.9 | 79.4 | 77.7 | 78.5 | 78.5 |
| DPASRF | Glass Brain | 78.8 | 79.8 | 79.3 | 80.6 | 79.7 | 80.2 | 79.9 | 80.7 | 79.8 | 79.9 | 79.8 | 79.8 | 78.8 | 79.1 | 78.9 | 78.8 | 79 | **81.3** | 80.1 | 81.4 |
|  | Stat_Map | 78.9 | 78.9 | 78.9 | 81.2 | 77.8 | 79.7 | 78.7 | 79.8 | 78.6 | 78.9 | 78.7 | 78.9 | 77.8 | 79.9 | 78.8 | 80.4 | 80.1 | 78.2 | 79.1 | 81.6 |
| NIAK | Glass Brain | 79.8 | 80 | 79.9 | **81.5** | 79.6 | 80.1 | 79.4 | **82.1** | 79.7 | 79.8 | 79.7 | 80.2 | 80.2 | 78.5 | 79.3 | 80.6 | 77.8 | 81 | **80.9** | 80.6 |
|  | Stat_Map | 76.9 | 79.6 | 78.3 | 78.8 | 78.7 | 80.3 | 79.5 | 80.3 | 77.9 | 78.6 | 78.3 | 79.6 | 79.8 | 77.8 | 78.8 | 78.7 | 76.9 | 79.4 | 78.1 | 79.6 |

The maximum performance is marked as bold in each section. The highest average accuracy obtain in our proposed method for CPAC glass brain images is 83%, average precision 80.5% for CCS stat_map images, average F1-score 80.9% for DPARSF glass brain images and average specificity 81.2% for NIAK glass brain images. From the analysis of the performance comparison, it shows that the performance of the improved CNN over other methods is statistically noteworthy.

### E. Deep Ensemble Classifier Performances Analysis

We perform four ensemble classifier techniques in the experiment by combining benchmark approaches with improved CNN. The ensemble classifiers are defined as ensemble classifier 1 (DenseNet + Proposed CNN), ensemble classifier 2 (ResNet + Proposed CNN), ensemble classifier 3 (Xception + Proposed CNN) and ensemble classifier 4 (Inception V3 + Proposed CNN), respectively.

All the ensemble classifier trained using ADAM optimizer and sigmoid function as classification. The final output was taken based on the equation (6) for the binary classification. Table V represents the classification performance for ensemble learning showing that the third ensemble classifier performs better than other ensembles. The highest accuracy and other relevant measurements are marked as bold in the table for each pipeline generated images.

**Table V:** Deep Ensemble learning classifier performance comparison.

| Pipelines | Ensemble Classifier 1 |  |  |  | Ensemble Classifier 2 |  |  |  | Ensemble Classifier 3 |  |  |  | Ensemble Classifier 4 |  |  |  |
| --- | --- | --- | --- | --- | --- | --- | --- | --- | --- | --- | --- | --- | --- | --- | --- | --- |
|  | P | R | F | A | P | R | F | A | P | R | F | A | P | R | F | A |
| CCS | 82.3 | 79.9 | 81.1 | 86 | 85.6 | 84.5 | 85.1 | 86 | **86.8** | 85.2 | 86 | **87** | 86.5 | **85.8** | **86.2** | 84 |
| CPAC | 83.2 | 81.2 | 82.2 | 85 | **86.4** | 85.3 | **85.8** | 85 | 85.2 | 83.8 | 84.5 | **87** | 82.4 | **86.4** | 84.4 | 85 |
| DPARF | 86.1 | 82.3 | 84.2 | 85 | 82.4 | 82.3 | 82.4 | 86 | **86.7** | **85.9** | **86.3** | **86** | 83.5 | 84.1 | 83.8 | 85 |
| NIAK | 83 | 84.1 | 83.5 | 86 | 81.9 | 83.4 | 82.6 | **86** | **86.8** | 81.8 | 84.2 | 85 | 84.6 | 84.5 | **84.5** | 84 |

### F. Unit Site Classification Performances Analysis

As the ABIDE is a consortium of the large dataset for ASD subjects from multiple renowned institutions around the world, therefore we performed unit-site classification to pattern out the site’s variability. Table VI represents the performance comparison using the proposed CNN model, including benchmark approaches. In our experiments, the highest accuracy for glass brain images is up to 85% belongs to several data sites and for stat_map images the accuracy is up to 84% for several data sites as shown in the table. Heinsfeld et al. also investigated the unit-site classification, where they achieved the highest accuracy of 68%, both for Caltech and MaxMun data site by using patterns of functional connectivity [23]. The results comparing with other literature suggest that there has data variability (dispersion or spread) among these sites that do not exist in other sites.

**Table VI:** Unit site classification accuracy comparison.

| Site-Out | Generated Images | DenseNet (%) | ResNet (%) | Xception (%) | Inception V3 (%) | Proposed Method (%) |
| --- | --- | --- | --- | --- | --- | --- |
| Caltech | Glass Brain | 81 | 79 | 80 | 82 | 84 |
|  | Stat_Map | 80 | 80 | 81 | 83 | 82 |
| CMU | Glass Brain | 79 | 79 | 80 | 79 | 83 |
|  | Stat_Map | 80 | 78 | 78 | 80 | **84** |
| KKI | Glass Brain | 80 | 81 | 79 | 80 | 83 |
|  | Stat_Map | 78 | 78 | 78 | 78 | 82 |
| Leuven | Glass Brain | 82 | 83 | 81 | 82 | 84 |
|  | Stat_Map | 80 | 80 | 80 | 80 | 84 |
| MaxMun | Glass Brain | 79 | 79 | 79 | 80 | 82 |
|  | Stat_Map | 78 | 80 | 82 | 80 | 81 |
| NYU | Glass Brain | 84 | 82 | 83 | 84 | **85** |
|  | Stat_Map | 80 | 81 | 82 | 79 | 83 |
| OHSU | Glass Brain | 79 | 80 | 81 | 78 | 82 |
|  | Stat_Map | 77 | 78 | 82 | 77 | 84 |
| OLIN | Glass Brain | 82 | 83 | 80 | 82 | 82 |
|  | Stat_Map | 79 | 82 | 78 | 82 | 83 |
| PITT | Glass Brain | 80 | 82 | 84 | 80 | 84 |
|  | Stat_Map | 79 | 82 | 83 | 79 | 81 |
| SBL | Glass Brain | 77 | 77 | 78 | 77 | 82 |
|  | Stat_Map | 77 | 78 | 78 | 77 | 80 |
| SDSU | Glass Brain | 78 | 79 | 79 | 78 | 79 |
|  | Stat_Map | 78 | 78 | 79 | 78 | 80 |
| Stanford | Glass Brain | 79 | 80 | 78 | 78 | 79 |
|  | Stat_Map | 78 | 80 | 80 | 79 | 80 |
| Trinity | Glass Brain | 80 | 80 | 81 | 83 | 84 |
|  | Stat_Map | 80 | 81 | 83 | 80 | 84 |
| UCLA | Glass Brain | 82 | 83 | 82 | 81 | 84 |
|  | Stat_Map | 83 | 81 | 82 | 83 | 84 |
| UM | Glass Brain | 84 | 83 | 84 | 84 | **85** |
|  | Stat_Map | 84 | 84 | 83 | 84 | 83 |
| USM | Glass Brain | 82 | 83 | 84 | 84 | 84 |
|  | Stat_Map | 79 | 82 | 81 | 84 | 84 |
| Yale | Glass Brain | 82 | 83 | 80 | 80 | 83 |
|  | Stat_Map | 83 | 80 | 80 | 79 | 83 |
| Mean |  | **80.1** | **80.6** | **80.7** | **80.4** | **82.7** |

## V. CONCLUSION

The recent advancement of functional connectivity and brain ROIs analysis has made a conspicuous invasion into the following classification of ASD. However, it is challenging to generalize the outcomes for larger, more heterogeneous populations rather than for smaller ones. While most of the recent works investigated the functional connectivity or time series analysis of fMRI, in this study, we demonstrate a suitable image generator to harvest a stable image that can provide perceptive details on the target disease using heterogeneous neuroimaging modality. Also, we validate the generated images using two propose deep learning-based frameworks that could enhance diagnostic truthfulness, with the potential to classify and develop better treatments. Our significance image processing scheme and sampling along with the precise CNN classifier ensure trustworthy approach of ASD classification associating with the other image processing techniques. Overall, the proposed image processing scheme provides a proficient and objective way of interpreting neuroimaging applied to the deep learning model.

Future research inclinations involve expending structural preprocessing and calculation of cortical measures pipeline data for classification of ASD. Besides, it is necessary to modify the architecture to consolidate raw fMRI data as well as to analyze the correlation between brain activation regions (axial, sagittal and coronal) to perceive the neural connectivity of brain during the natural progression of autism.

## ACKNOWLEDGMENT

This work was supported in part by the National Natural Science Foundation of China under Grant 61572231, and in part by the Shandong Provincial Key Research & Development Project Grant 2017GGX10141.

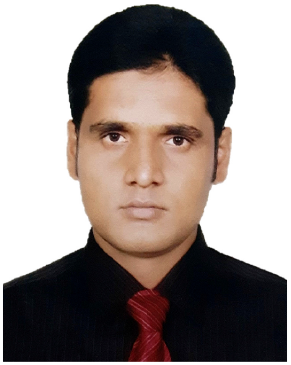

**Md Rishad Ahmed** is a Chinese Government Scholarship holder with Shandong Provincial Key Laboratory of Network Based Intelligent Computing, University of Jinan, China where he is currently a Computer Science Masters Degree student. He received his M.Sc. and B.Sc. in Applied Physics and Electronic Engineering from University of Rajshahi, Bangladesh. His research interests are biomedical image processing, machine learning, deep learning, pattern recognition, and classification.

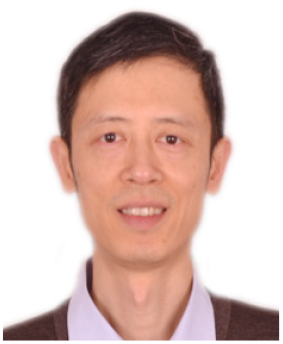

**Yuan Zhang** received his M.S. degree in Communication Systems and Ph.D. degree in Control Theory & Engineering (Biomedical Engineering) both from Shandong University, China, in 2003 and 2012 respectively. He is currently an Associate Professor at University of Jinan, China. Dr. Zhang was a visiting professor at Computer Science Department, Georgia State University, USA, in 2014. As the first author or corresponding author he has published more than 50 peer reviewed papers in international journals and conference proceedings, 1 book chapters, and 6 patents in the areas of Smart Health and Biomedical Big Data Analytics. He has served as Leading Guest Editor for six special issues of IEEE, Elsevier, Springer and InderScience publications, including IEEE Internet of Things Journal special issue on Wearable Sensor Based Big Data Analysis for Smart Health and IEEE Journal of Biomedical and Health Informatics (JBHI) special issue on Pervasive Sensing and Machine Learning for Mental Health. He has served on the technical program committee for numerous international conferences. He is an associate editor for IEEE Access and Internet of Things (Elsevier). Dr. Zhang’s research interests are Wearable Sensing for Smart Health, Machine Learning for Auxiliary Diagnosis, and Biomedical Big Data Analytics. His research has been extensively supported by the Natural Science Foundation of China, China Postdoctoral Science Foundation, and Natural Science Foundation of Shandong Province with total grant funding over 1.5 million RMB. Dr. Zhang is a Senior Member of both IEEE and ACM. For more information, please refer to http://uslab.ujn.edu.cn/index.html.

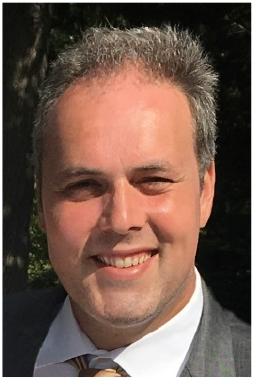

**Dr. Omer T. Inan** (S’06, M’09, SM’15) received the B.S., M.S., and Ph.D. degrees in electrical engineering from Stanford University, in 2004, 2005, and 2009, respectively. From 2009-2013, he was a Visiting Scholar in the Department of Electrical Engineering, Stanford University. Since 2013, Dr. Inan has been a faculty member at the Georgia Institute of Technology where he is currently Associate Professor of Electrical and Computer Engineering. His research focuses on non-invasive physiologic sensing and modulation for human health and performance. Dr. Inan is an Associate Editor of the IEEE Journal of Biomedical and Health Informatics, Associate Editor for the IEEE Engineering in Medicine and Biology Conference and the IEEE Biomedical and Health Informatics Conference, and Technical Program Committee Member or Track Chair for several other major international biomedical engineering conferences. He has published more than 125 technical articles in peer reviewed international journals and conferences, and has multiple issued patents. Dr. Inan received the Gerald J. Lieberman Fellowship in 2009, the Lockheed Dean’s Excellence in Teaching Award in 2016, the Sigma Xi Young Faculty Award in 2017, the IEEE Sensors Early Career Award in 2018, the Office of Naval Research Young Investigator Award in 2018, and the National Science Foundation CAREER Award in 2018.

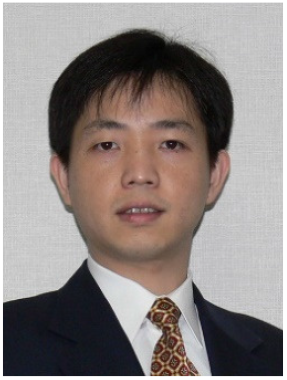

**Hongen Liao** (M’04, SM’14) received the B.S. degree in mechanics and engineering sciences from Peking University, Beijing, China, in 1996, and the M.E. and Ph.D. degrees in precision machinery engineering from the University of Tokyo, Tokyo, Japan, in 2000 and 2003, respectively. He was a Research Fellow of the Japan Society for the Promotion of Science. Since 2004, he has been a faculty member at the Graduate School of Engineering, University of Tokyo, where he became an Associate Professor in 2007. He has been selected as a National “Thousand Talents” Distinguished Professor, National Recruitment Program of Global Experts, China, and is currently a full Professor and Vice Director in the Department of Biomedical Engineering, School of Medicine, Tsinghua University, Beijing, China. His research interests include 3-D medical image, image-guided surgery, medical robotics, computer-assisted surgery, and fusion of these techniques for minimally invasive precision diagnosis and therapy. He is the author and co-author of more than 250 peer-reviewed articles and proceedings papers, as well as 40 patents, over 290 abstracts and numerous invited lectures. He is an Associate Editor of IEEE Engineering in Medicine and Biology Society Conference, the Organization Chair of Medical Imaging and Augmented Reality Conference (MIAR) 2008, the Program Chair of the Asian Conference on Computer-Aided Surgery Conference (ACCAS) 2008 and 2009, the Tutorial co-chair of the Medical Image Computing and Computer Assisted Intervention Conference (MICCAI) 2009, the Publicity Chair of MICCAI 2010, the General Chair of MIAR 2010 and ACCAS 2012, the Program Chair of MIAR 2013, the Workshop Chair of MICCAI 2013 and 2019, and the General co-chair of MIAR 2016. He is a President of Asian Society for Computer Aided Surgery and Co-chair of Asian-Pacific Activities Working Group, Inter-national Federation for Medical and Biological Engineering (IFMBE).

## Footnotes

1 http://preprocessed-connectomes-project.org/abide/download.html

